# A structure-guided approach to predict MHC-I restriction of T cell receptors for public antigens

**DOI:** 10.1101/2024.06.04.597418

**Authors:** Sagar Gupta, Nikolaos G. Sgourakis

**Affiliations:** Center for Computational and Genomic Medicine, Department of Pathology and Laboratory Medicine, Children’s Hospital of Philadelphia, Philadelphia, PA, USA; Department of Biochemistry and Biophysics, Perelman School of Medicine at the University of Pennsylvania, Philadelphia, PA, USA

## Abstract

Peptides presented by class I major histocompatibility complex (MHC-I) proteins provide biomarkers for therapeutic targeting using T cell receptors (TCRs), TCR-mimicking antibodies (TMAs), or other engineered protein binders. Despite the extreme sequence diversity of the Human Leucocyte Antigen (HLA, the human MHC), a given TCR or TMA is restricted to recognize epitopic peptides in the context of a limited set of different HLA allotypes. Here, guided by our analysis of 96 TCR:pHLA complex structures in the Protein Data Bank (PDB), we identify TCR contact residues and classify 148 common HLA allotypes into T-cell cross-reactivity groups (T-CREGs) on the basis of their interaction surface features. Insights from our work have actionable value for resolving MHC-I restriction of TCRs, guiding therapeutic expansion of existing therapies, and informing the selection of peptide targets for forthcoming immunotherapy modalities.

## Introduction

The class I major histocompatibility complex (MHC-I) molecules display endogenous peptides that are recognized by CD8^+^ T cell receptors (TCRs) to allow for identification and elimination of infected or cancerous cells^1^. Following the first structure of an αβ TCR recognizing its peptide/MHC-I target^2^, a large number of co-crystal structures have revealed the molecular basis of TCR antigen recognition^3^. Notwithstanding, the Human Leukocyte Antigen class I (HLA-I) locus^4^ encodes for more than 35,000 different protein allotypes within the classical (*HLA-A*, *HLA-B*, and *HLA-C*) and nonclassical (*HLA-E*, *HLA-F*, and *HLA-G*) genes^5^. Each HLA allotype can present a distinct repertoire of at least 10^8^ epitopic peptides as peptide/HLA (pHLA) complexes, ensuring a wide coverage of self or aberrant peptide repertoires at the population scale^6^. HLA peptide binding preferences form the basis for their classification into supertypes that have been widely adopted by the immunology community^7,8^. On the other hand, TCRs are restricted to recognize peptides in the context of specific HLA allotypes using their germline-encoded complementarity-determining regions 1 and 2 (CDR1 and −2) loops which primarily contact the helical regions of the MHC^9^. Specific recognition of epitopic peptides by TCRs occurs primarily through the highly variable CDR3 loops, which are generated via somatic recombination of the variable (V), joining (J), and, in β chains, diversity (D) genes^3^. Thus, despite the lack of a precise recognition “code”, a diverse repertoire of 10^8^ TCRs that are present in any individual donor^10^ likely employ a limited set of possible configurations, or docking angles, to engage their pHLA targets^3,9,11,12^.

While the exact model for MHC restriction of TCR recognition has been debated (reviewed in ref. ^13^), it is clear that TCRs can be alloreactive i.e., recognize the same peptide antigen in the context of different HLA allotypes^13–20^. A notable example is the LC13 TCR that reacts with peptide-bound HLA-B*08:01 and HLA-B*44:05^15^. Structural studies revealed that a specific TCR binding mode enabled avoidance of surface polymorphisms between these two HLA-B allotypes, in contrast to two other non-alloreactive TCRs (RL42 and CF34) derived from HLA-B8^+^-B44^+^ individuals^21^. Furthermore, in our recent characterization of a TCR-mimicking antibody (TMA), 10LH, we showed that this scFv was restricted to recognize a public neuroblastoma epitope derived from the PHOX2B oncoprotein in the context of HLAs belonging to the A9 serological cross-reactivity group (CREG)^22^. These results suggest parallels in HLA restriction profiles between TCRs and HLA cross-reactivity of alloreactive antibodies against donor HLA molecules, as established in the organ transplantation community over two decades ago^23^.

Here, by analyzing 96 TCR:pHLA complex structures from the Protein Data Bank (PDB)^24^, we uncover that the level of polymorphisms at TCR contact residues is dependent on the HLA allotype. Leveraging this observation, we create an unbiased classification system for HLA allotypes, called T-cell cross-reactivity groups (T-CREGs), to cluster 148 common HLAs based on their sequence similarity of TCR contact residues. We apply T-CREGs to characterize the TCR cross-reactivity landscape of public epitopes and find that information gleaned from our analysis can be used to predict the HLA restriction of known TCRs, rank emerging epitopes according to the number of receptors required to maximize patient coverage, and guide patient cohort expansion for existing TCR or antibody-based therapeutics.

## Results

To evaluate the extent of HLA polymorphisms of TCR-contacting residues, we mined TCR3d^25^ and identified 96 TCR:pHLA crystal structures. Across all ternary complexes, we identified a set of 32 TCR contacts, defined as MHC-I residues with a side chain or Cα atom within 5 Å of any TCR atom in at least 10% of structures^26^. As expected, the TCR-contacting residues spanned the solvent-exposed positions on the α_1_ and α_2_ helices on the MHC-I structure, including the previously identified restriction triad^27^. To visualize polymorphisms across these positions, we created sequence logos for common HLA alleles (>0.05% American allelic frequency)^28^. We observed that TCR-contacting residues were more polymorphic in HLA-A* alleles in comparison to HLA-B* and HLA-C* alleles (**Fig. 1**). Notably, MHC-I peptide-binding groove residues did not follow this allotype-specific trend and showed a similar level of divergence across HLA loci (**Extended Data Fig. 1**). Our observed sequence-based trends suggested that different HLA allotypes share conserved molecular surfaces that can be used as a basis for understanding and predicting their TCR restriction.

**Figure 1.**
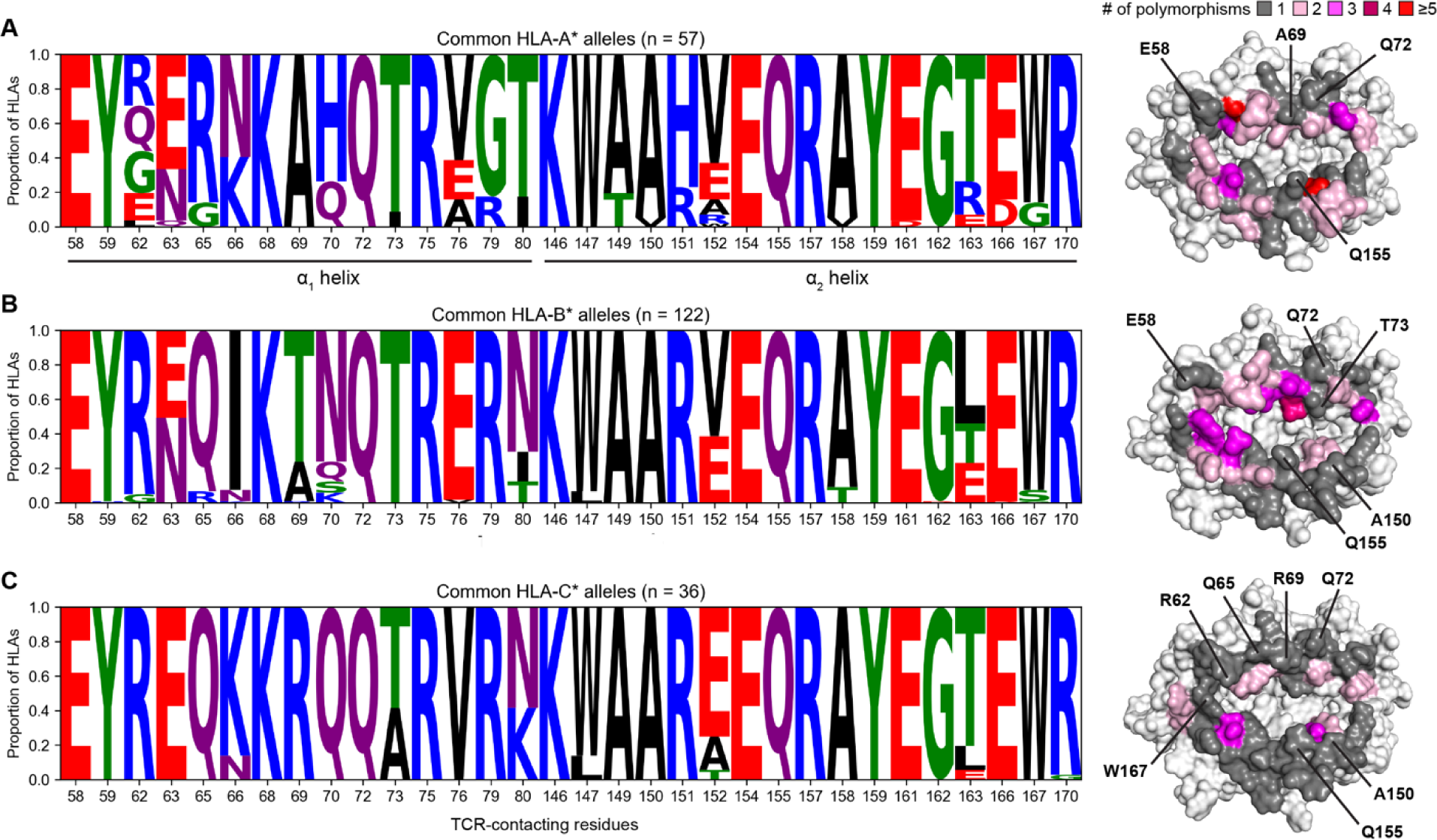
Residue polymorphisms of TCR interaction sites are HLA locus dependent. **(A** to **C).** Sequence logo depicting the amino acid identity at a given TCR-contacting residue across a total of 215 common (A) HLA-A*, (B) HLA-B*, and (C) HLA-C* alleles. To the right, sequence conservation across common HLA alleles is mapped onto the (A) HLA-A*02:01 surface (PDB ID 1DUZ), (B) HLA-B*07:02 surface (PDB ID 4U1H), (C) HLA-C*06:02 surface (PDB ID 5W6A). The MHC-I surface is colored according to the number of polymorphisms present at each position as derived from the sequence logo. Non-TCR-contacting residues are colored white. Select, conserved TCR-contacting residues are marked on each HLA surface.

To further classify HLA molecular surfaces based on of their specific TCR recognition mode, we clustered HLA alleles according to the amino acid similarity of TCR-contacting residues (**Fig. 2A**). Recognizing that identical molecular surfaces are not a strict requirement for TCR binding^15,22^, we established the HLA similarity criterion based on biochemical identity and BLOSUM62 score^29^. If two HLAs satisfy this criterion, they are called neighbors. Then, we applied a greedy algorithm to determine a minimal set of HLA alleles which could capture the diversity of molecular surface among common HLAs. Briefly, a binary symmetric adjacency matrix was constructed based on our HLA similarity criterion. The HLA allele with the most neighbors was assigned to a T-CREG and, along with its neighboring alleles, was removed from the matrix. This procedure was continued iteratively until all HLAs were covered. Each T-CREG is named according to the allele with the highest allelic frequency assigned to it. Using this ansatz, we found that our set of 215 common HLA alleles could be reduced to a minimal set of 62 T-cell cross-reactivity groups, or T-CREGs. By filtering T-CREGs based on their American allelic frequency coverage^28^, we established a final list of 21 T-CREGs that include 148 alleles (**Table 1**). This list covers 74.1%, 86.2%, and 89.5% of the US population according to HLA-A*, HLA-B*, and HLA-C* haplotypes, respectively (**Fig. 2B**). Despite our use of an unbiased approach, each T-CREG contained HLAs of the same allotype, except in the case of the C*03:04 T-CREG which contained B*46:01, an allele which has been reported to arise from intergenic mini-conversion between B*15:01 and C*01:02^30^. We also capture the C1/C2 dimorphism by separating C*05:01 and C*08:02 into different T-CREGs^31^. Notably, the ratio of HLA-A* alleles to T-CREGs is disproportionately lower at 3.7 alleles per T-CREG in comparison to HLA-B* (10.7 alleles per T-CREGs) and HLA-C* (5.0 alleles per T-CREG) alleles. This agrees with our prior finding that HLA-A* alleles have more polymorphisms among their TCR-contacting residues (**Fig. 1**). To quantify and visualize the diversity of HLA molecular surfaces, we used a two-dimensional PCA plot based on the BLOSUM62 score of individual TCR-contacting residues (**Fig. 2C**). In agreement with our T-CREG analysis, we find that HLA-A* alleles were more dispersed in the PCA plane compared to HLA-B* and HLA-C* alleles.

**Figure 2.**
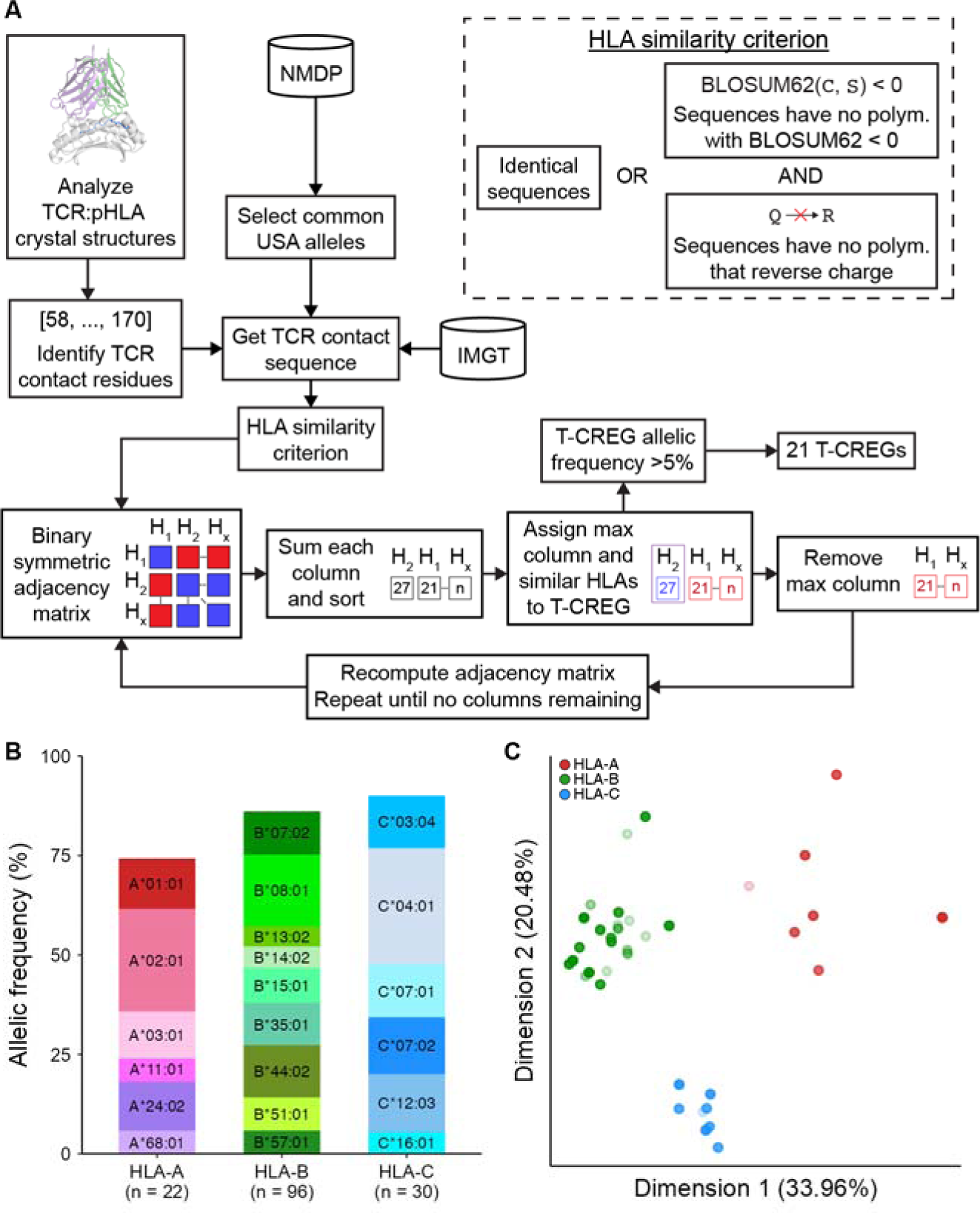
Structure-guided classification of HLA allotypes into T-CREGs. **(A)** Schematic depicting the selection of T-CREGs. Briefly, the TCR contact sequence is acquired by identifying TCR-contacting residues across common alleles via the NMDP database and obtaining sequences from the IMGT database. Then, using the HLA similarity criterion, the T-CREGs are selected using a greedy algorithm. **(B)** Stacked bar plot showing the 21 T-CREGs and the proportion of allelic frequency they cover. The number of HLA alleles included in the T-CREGs of a given allotype is shown in the x-axis. **(C)** A two-dimensional PCA plot based on the BLOSUM62 score of the 32 TCR-contacting residues explains 54.44% of the variance. Each point corresponds to one of the 148 HLAs covered by the 21 T-CREGs and is colored according to the allotype. Individual data points can be found in the Source Data file.

**Table 1.** T-CREGs with a combined USA allelic frequency greater than 5%.

| **T-CREG name** | **Combined USA allelic frequency** | **TCR-contacting residue polymorphisms** | **T-CREG members** |
| --- | --- | --- | --- |
| A*01:01 | 12.29% | None | A*01:01, A*01:02 |
| A*02:01 | 26.21% | None | A*02:01, A*02:02, A*02:04, A*02:05, A*02:06, A*02:07, A*02:10, A*02:17, A*02:22, A*02:60 |
| A*03:01 | 11.39% | None | A*03:01 |
| A*11:01 | 5.86% | None | A*11:01, A*11:02 |
| A*24:02 | 12.72% | R151H | A*23:01, A*24:02, A*24:17, A*24:25 |
| A*68:01 | 5.61% | None | A*68:01, A*68:02, A*69:01 |
| B*07:02 | 10.45% | N63E, A69T, Q70N | B*07:02, B*07:05, B*15:40 |
| B*08:01 | 18.45% | N63E, T69A, N70Q, N80T, A158T | B*08:01, B*18:01, B*18:02, B*37:01, B*38:02, B*39:01, B*39:02, B*39:05, B*39:06, B*39:08, B*39:09, B*39:10, B*39:11, B*41:01, B*41:02, B*41:03, B*42:01, B*42:02, B*54:01, B*55:02, B*67:01 |
| B*13:02 | 5.05% | E63N, T80N | B*13:01, B*13:02, B*40:02, B*40:03, B*40:04, B*40:06, B*40:08, B*40:11, B*40:27, B*47:01 |
| B*14:02 | 5.02% | T69A, N70Q | B*14:01, B*14:02, B*14:03, B*55:01 |
| B*15:01 | 9.16% | E63N, T69A, N70Q | B*15:01, B*15:02, B*15:03, B*15:04, B*15:07, B*15:08, B*15:09, B*15:10, B*15:11, B*15:15, B*15:150, B*15:18, B*15:21, B*15:25, B*15:27, B*15:29, B*15:30, B*15:35, B*15:37, B*15:39, B*35:14, B*35:43, B*40:05, B*50:01, B*56:03, B*78:01 |
| B*35:01 | 10.67% | E63N, T69A, N70Q | B*15:05, B*35:01, B*35:02, B*35:03, B*35:04, B*35:05, B*35:08, B*35:12, |
|  |  |  | B*35:17, B*35:20, B*48:02, B*56:01, B*56:04 |
| B*51:01 | 8.75% | N63E | B*15:13, B*49:01, B*51:01, B*51:02, B*51:06, B*51:07, B*52:01 |
| B*44:02 | 12.73% | T80N | B*44:02, B*44:03, B*44:05, B*44:10, B*45:01 |
| B*57:01 | 5.63% | None | B*57:01, B*57:02, B*57:03, B*57:04, B*58:01, B*58:02 |
| C*03:04 | 12.78% | None | B*46:01, C*03:02, C*03:03, C*03:04, C*03:05, C*03:06 |
| C*12:03 | 15.00% | T73A | C*01:02, C*01:03, C*08:02, C*08:04, C*12:02, C*12:03, C*14:02, C*14:03 |
| C*04:01 | 29.54% | A73T | C*04:01, C*04:03, C*04:04, C*04:06, C*04:07, C*05:01, C*06:02, C*12:04, C*18:01 |
| C*07:01 | 13.41% | None | C*07:01, C*07:26 |
| C*07:02 | 13.88% | None | C*07:02, C*07:04 |
| C*16:01 | 5.25% | T152A | C*08:01, C*08:03, C*16:01, C*16:04 |

To evaluate the accuracy of our T-CREG classification scheme in the absence of high-resolution TCR:pHLA structures for a given HLA of interest, we identified TCR-contacting residues for each HLA that is present in TCR3d, on a per-structure basis. Then, we used our HLA similarity criterion to determine which HLA surfaces could by potentially cross-reactive with the TCR present in each structure. We quantified the number of instances that a T-CREG could completely map to the set of structure-derived HLA alleles. For instance, if a TCR was identified to interact with only two out of three members of the A*68:01 T-CREG e.g., A*68:01 and A*68:02 but not A*69:01, then the T-CREG can be further “split”. This could occur if the TCR-contacting residues associated with a particular structure included a polymorphic residue not identified in our aggregate residue list, causing the T-CREG to be split. We found that the T-CREGs were complete in 92.7% of structures (**Extended Data Fig. 2**). Given that we employed a broad definition of TCR-contacting residues by including nearly every solvent-exposed residue on the α_1_ and α_2_ helices, this finding was expected. This result supports that T-CREGs can be broadly applicable to provide an initial assessment of HLA restriction for TCRs and TMAs, in the absence of experimental structures.

To further demonstrate possible applications of our work to support the development of autologous T cell or peptide-centric CAR-T therapies using TMAs^32^, we applied T-CREGs for all peptide epitopes in our dataset of TCR:pHLA crystal structures, as well as our recently solved structure^22^ of an TCR-mimicking antibody, 10LH, targeting the PHOX2B peptide (QYNPIRTTF) presented by A*24:02. For each peptide, we predicted the strong binders among common HLAs by NetMHCPan-4.1^33^ and identified distinct molecular surfaces using the T-CREGs. We highlight seven public epitopes which include established cancer antigens such as MAGE-A3 (EVDPIGHLY)^34^, PHOX2B (QYNPIRTTF)^35^, and gp100 (YLEPGPVTV)^36^ (**Fig. 3A**). As we have recently developed and characterized a TMA for PHOX2B/A*24:02^22,35^, we focus on this epitope. The 9mer peptide binds strongly to 35 common alleles which map to 9 T-CREGs including our previously identified A9 cluster of alleles (**Fig. 3B**). These results suggest that future efforts should be focused towards developing binders towards alleles in the C*04:01, C*07:02, and C*07:01 T-CREGs, which can maximize therapeutic expansion. Visual inspection of these alleles’ TCR-contacting residues and modeling of their molecular surfaces confirms that there are likely sufficient differences that would prohibit the existing TMA from engaging these PHOX2B/HLA-C* complexes (**Fig. 3C**). Thus, the T-CREGs are a useful tool for prioritizing tumor-associated epitopes and for identifying possible routes for expanding therapies using a minimal set of therapeutics.

**Figure 3.**
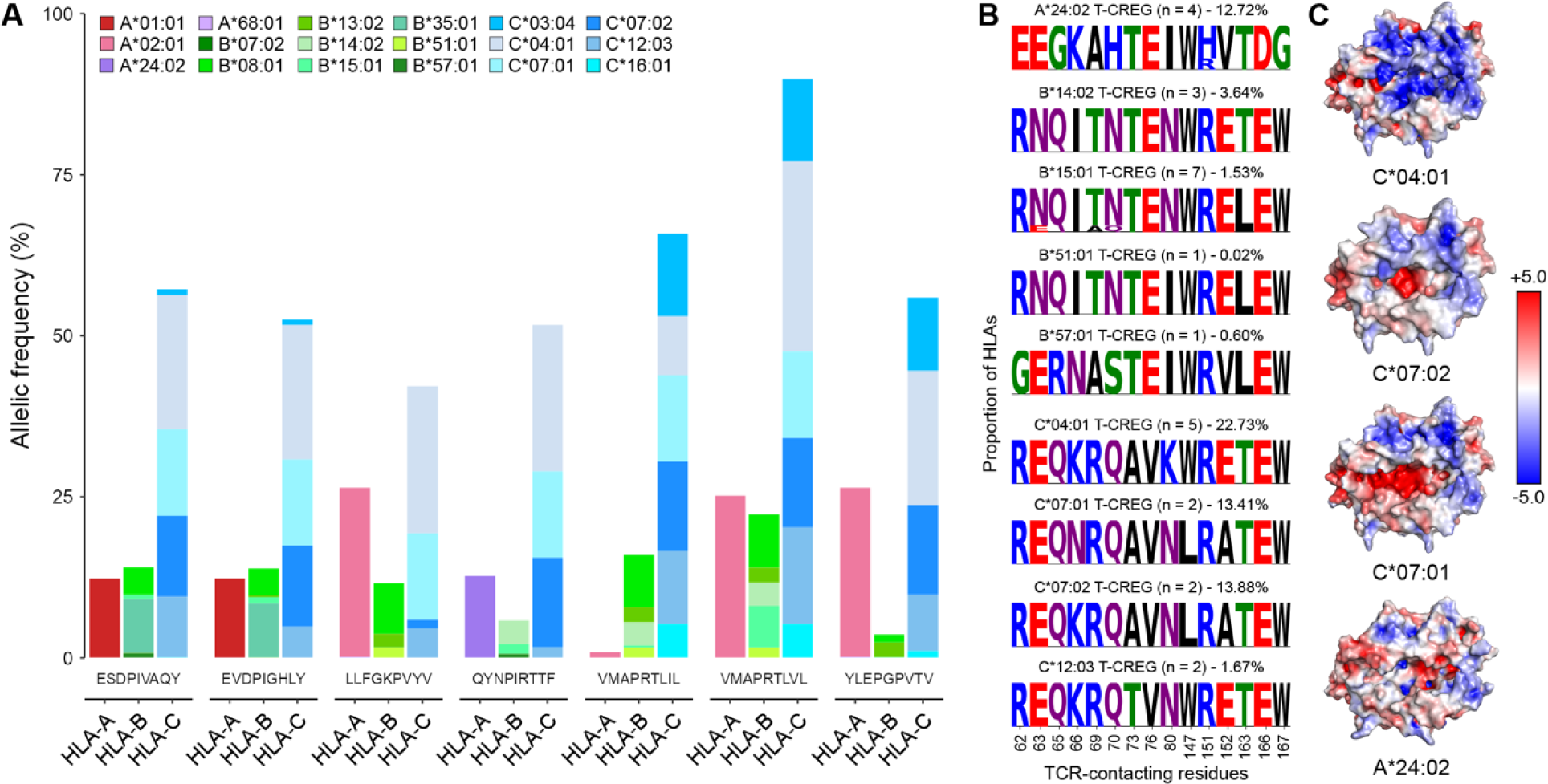
Application of T-CREGs to estimate therapeutic coverage of public epitopes. **(A)** Stacked bar plot showing the allelic frequency of HLAs that are predicted strong binders to peptide antigens in TCR3d, including PHOX2B (QYNPIRTTF). The HLAs were then classified into T-CREGs and split by allotype. Antigens were only shown if they crossed the 40% allelic frequency threshold in at least one allotype. **(B)** Sequence logos depicting the amino acid identity at a given non-conserved TCR-contacting residues across T-CREGs with at least one predicted strong PHOX2B-binding allele. The percentage indicates the allelic frequency of the American population covered by the predicted strong binding alleles in a given T-CREG. **(C)** Electrostatic representation of the molecular surfaces of PHOX2B-binding T-CREGs with the highest allelic frequency coverage. PDB ID 7MJA was used for A*24:02 and all other HLA-C* allele models were generated using RosettaRemodel and the FastRelax application. The HLA with the highest allelic frequency in each T-CREG is shown.

A limitation of using T-CREGs to infer TCR cross-reactivity among HLAs presenting the same peptide antigen is the assumption that the peptide conformation will be conserved. We wanted to understand if peptide conformational changes were being driven by the intrinsic flexibility of the peptide or by HLA groove polymorphisms. Thus, we measured structural differences between peptide backbones in complexes with the same pHLA sequence solved independently or in different asymmetric units extracted from a single pHLA crystal structure^37^. We find that in 31 out of 35 such cases, the peptide configuration was nearly identical, in agreement with a prior study^38^ (**Fig. 4A**). We also calculated structural differences between pHLA complexes with the same peptide sequence presented by different HLA allotypes and found significantly increased backbone deviations. To identify if there were specific polymorphisms prone to driving these structural changes, we compared the frequency of groove polymorphisms between cases where the peptide does and does not change conformation (**Fig. 4B**). As expected, polymorphisms were not found in HLA residues defining the A and F peptide binding pockets as changes here would impact antigen presentation^39^ (**Fig. 4C**). We discovered that polymorphisms at positions 116 and 142 had an increased association of cases with an altered peptide conformation. This supports that, if two alleles belonging to the same T-CREG are polymorphic at one of these positions, the peptide presented by the HLAs is likely to adopt a different configuration. As a result, separate therapeutics will be required to cover HLAs within the same T-CREG. These results support that, while T-CREGs can provide a robust lower-limit estimate for the number of therapeutics that are required to achieve a specific patient coverage, changes in peptide presentation among different HLAs should be considered explicitly using either experimental structure determination or computational modeling^40–47^.

**Figure 4.**
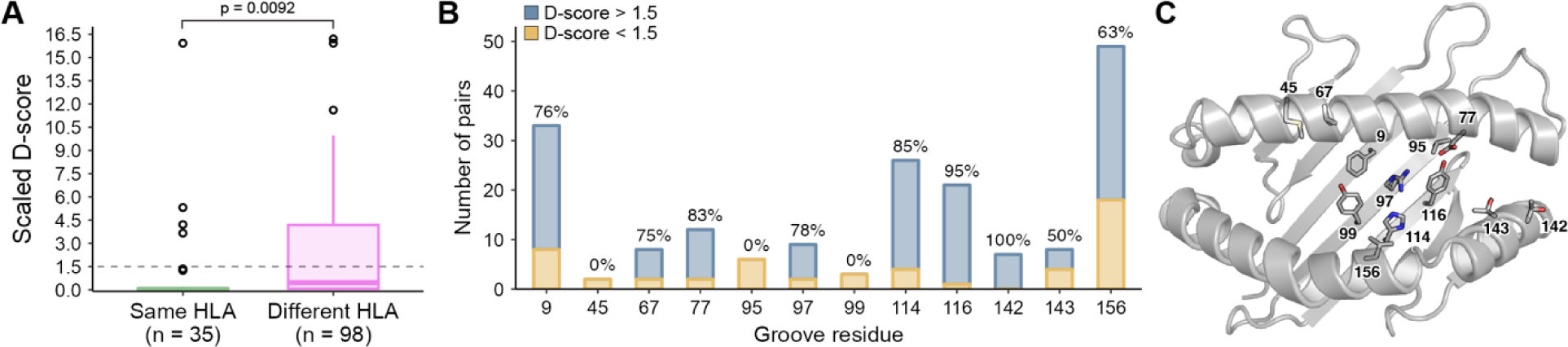
MHC-I groove polymorphisms impact antigen presentation by different HLAs. (**A**) Boxplot comparing the scaled D-scores of pHLA structures with the same peptide and same HLA sequence to those with the same peptide but different HLA sequence. Whiskers extend to the furthest values that lie within the 75th and 25th percentile value ± 1.5 times the interquartile range and outliers are shown in black circles. A two-sample unequal variance Welch’s t-test was performed to obtain the p-value of 0.0092. (**B**) Number of pairs that have at least one polymorphism at a given groove residue position. Residues with no polymorphisms are not shown. The percentage of pairs with a D-score greater than 1.5 is shown above each bar. (**C**) Relevant groove residues from panel (B) are shown as sticks on a cartoon representation of the HLA. PDB ID 1AO7 (A*02:01) was used for visualization.

## Discussion

Building upon our observation that conserved surface patterns bridge highly divergent HLA allotypes, we categorized HLAs based on their molecular surface features with respect to the TCR binding mode. Our work builds on the established serological cross-reactivity groups (CREGs) developed by the organ transplantation community^23^ and our prior findings on the structural principles that enabled a TCR-like antibody to recognize several HLAs^22^. We find that HLA-A* alleles are far more polymorphic and map to a disproportionately higher number of T-CREGs, relative to HLA-B* or HLA-C* allotypes. After validating the completeness of the T-CREGs, we identified possibilities to widen the range of binders targeting public epitopes.

We anticipate the T-CREGs being used in a variety of applications. For instance, if a public cancer neoantigen is identified^48^, researchers can prioritize the development of receptors against T-CREGs that cover a greater proportion of the patient population to maximize therapeutic benefit. Also, the number of patients eligible for existing TCR or TMA-based therapies could be expanded by rapidly testing for interactions with allotypes belonging to the same T-CREG as the original HLA which restricts a known receptor^22^. In the future, *de novo* receptor design^49–52^ could allow for binding modes that are directed towards conserved regions of the MHC-I surface, enabling cross-HLA targeting of public epitopes across different T-CREGs. Finally, our T-CREG analysis requires a minimal amount of information, simply the set of HLAs anticipated to bind the peptide, and is publicly available online through a code repository. In a manner similar to the classification of HLAs into supertypes on the basis of their peptide-binding preferences^7,8^, our work provides a toehold for understanding the HLA restriction of TCRs to guide experimental testing and therapeutic expansion in various research or clinical settings.

## Extended Data Figures

**Extended Data Figure 1.**
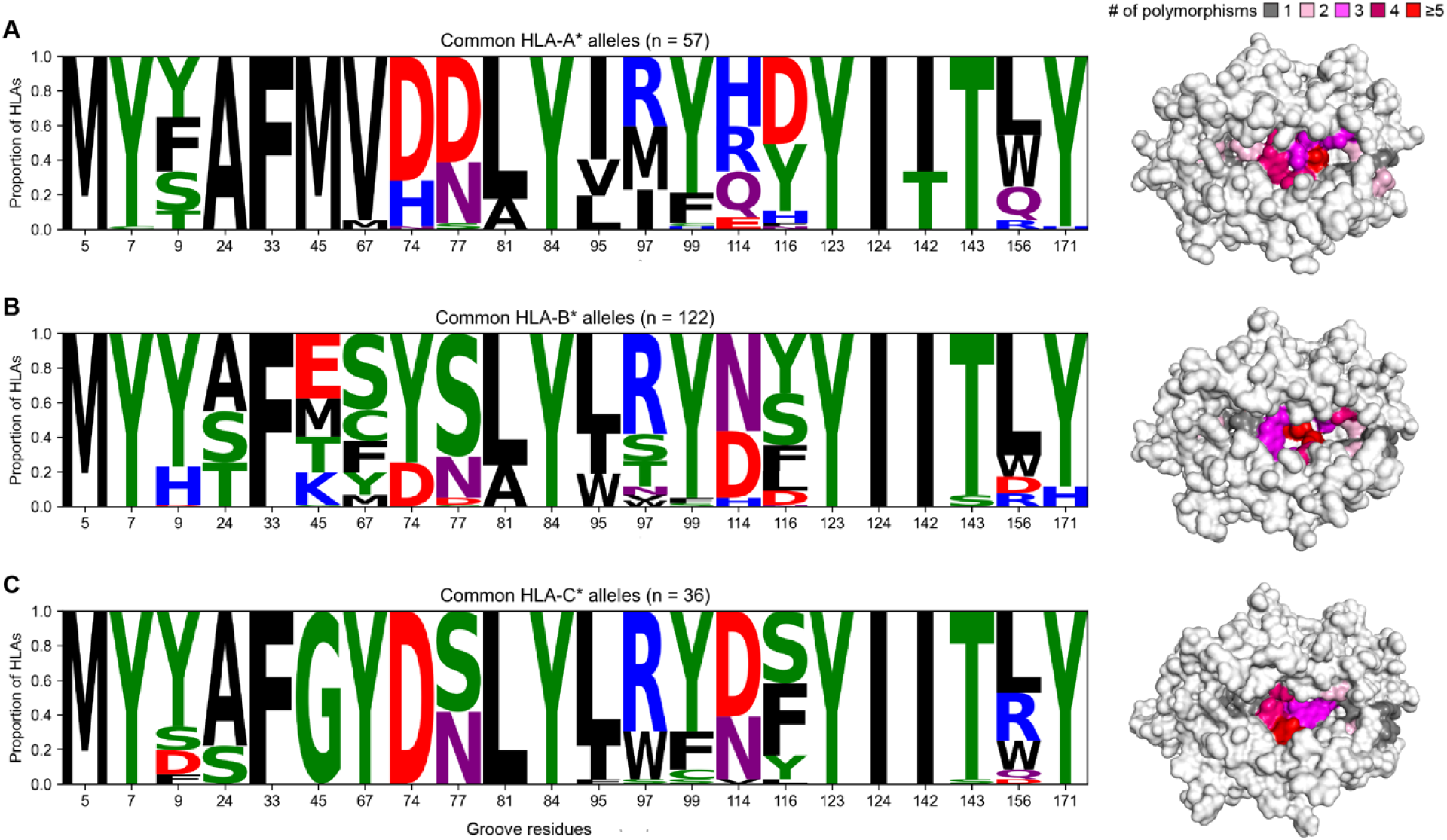
HLA polymorphisms of MHC-I peptide-binding groove residues. **(A** to **C).** Sequence logo depicting the amino acid identity at a given HLA groove residue across a total of 215 common (A) HLA-A*, (B) HLA-B*, and (C) HLA-C* alleles. To the right, sequence conservation across common HLA alleles is mapped onto the (A) HLA-A*02:01 surface (PDB ID 1DUZ), (B) HLA-B*07:02 surface (PDB ID 4U1H), (C) HLA-C*06:02 surface (PDB ID 5W6A). The MHC-I surface is colored according to the number of polymorphisms present at each position as derived from the sequence logo. Non-groove residues are colored white.

**Extended Data Figure 2.**
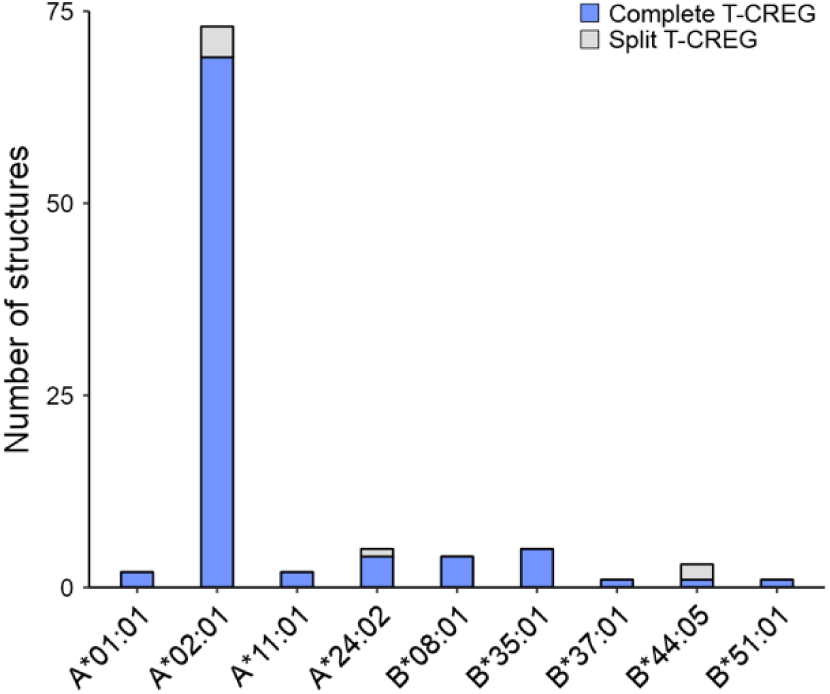
Validation of T-CREGs on a per-structure basis. The x-axis depicts the HLA allele of the analyzed structure.

## Methods

### Databases

Allelic (haploid) frequencies for each of the five major American race groups (African American, Asian or Pacific Islander, Caucasian, Hispanic, and Native American) were obtained from the National Marrow Donor Program (NMDP) database^28^. A combined American allelic frequency for each allele was obtained by normalizing values according to the racial distribution followed by the 2020 US Census. For alleles with an allelic frequency greater than 0.05%, HLA sequences were obtained from the IMGT/HLA database^5^, and a pairwise sequence alignment was performed to the HLA-A*02:01 heavy chain sequence (consisting of 180 residues from the N terminus Gly residue).

A total of 96 TCR:pHLA structures were obtained from TCR3d^25^ and 510 pHLA structures were obtained from HLA3DB^40^. Only structures with peptide lengths from 8 to 10 amino acids were included in our analysis.

### TCR contact and groove residue identification

For each TCR:pHLA structure, an HLA residue was a TCR-contacting residue if it had a side chain or Cα atom within 5 Å of a TCR atom after hydrogen atoms were removed from the structure using PyMOL (v. 2.5.3)^53^. Then, the set of TCR-contacting residues were determined by selecting residues that were identified as a TCR-contacting residue in at least 10% of structures^26^. Groove residues were identified using a similar criterion, where the distance was determined between the peptide and the HLA. If a groove residue was also a TCR-contacting residue, then it was not considered a groove residue. Sequence logos were created using the logomaker package in Python (v. 3.8.15).

### T-CREG identification

T-CREGs were identified using a greedy algorithm in Python (v. 3.8.15). A binary symmetric adjacency matrix was generated by determining the HLA similarity criterion for all pairs of HLAs. Then, for each HLA, the number of similar HLAs were summed. The allele with the greatest number of neighbors was noted, and it and its neighbors were removed from the matrix. This process continued iteratively until the matrix was empty. Then, the final T-CREGs list was determined by including T-CREGs with a combined allelic frequency of at least 5%.

### Analysis of public epitopes

NetMHCPan-4.1^33^ was used to determine strong (≤0.5% rank) binding alleles for each peptide in the TCR:pHLA dataset. Then, for each peptide, the T-CREGs were determined using the strong binding alleles as the input.

### Structural modeling

To model different alleles onto the PHOX2B/HLA-A*24:02 structure, we used the RosettaRemodel application^54^ from the Rosetta (v2020.08)^55^ suite of programs. A blueprint file was used to specify the polymorphisms relative to the HLA-A*24:02 sequence. The remodeled structures were then optimized using the FastRelax application^56^ three times and the optimized, remodeled structure with the lowest total Rosetta energy was selected as the final structure.

### Analysis of pairs of pHLA structures with the same peptide sequence

Peptide/HLA structures were obtained from HLA3DB^40^ and for each pair of structures with the same peptide sequence, the scaled D-score was calculated as described previously^40^. A two-sample unequal variance Welch’s t-test was conducted to compare the D-scores of the cases of same HLA and different HLA structures and significance was assessed according to a p-value cutoff of 0.01.

## Acknowledgments

The authors acknowledge Dr. Omar Ani for helpful discussions. This work was supported by NIH grants R01AI143997 (N.G.S.), R35GM125034 (N.G.S.), and The Children’s Hospital of Philadelphia Cell and Gene Therapy Collaborative. This work was delivered as part of the NexTGen and Matchmakers Teams supported by the Cancer Grand Challenges partnership funded by Cancer Research UK (CGCATF-2021/100002) and the National Cancer Institute (CA278687-01), and the Mark Foundation.

## Data and code availability

All data are available in the main text, in the supplementary materials, or provided in the Source Data file. Code used to determine the T-CREGs can be accessed on GitHub via: https://github.com/titaniumsg/find_tcreg/.

## Contributions

N.G.S. and S.G. conceived and designed the project. S.G. performed all methodology, investigation, and visualization. S.G. and N.G.S. wrote the paper, with feedback from all authors. N.G.S. acquired funding and supervised the project.

## Ethics declarations

All authors declare that the research was conducted in the absence of any commercial or financial relationships that could be construed as a potential conflict of interest.

